# Dietary nitrate and nitrite protect against doxorubicin-induced cardiac fibrosis and oxidative protein damage in tumor-bearing mice

**DOI:** 10.1101/2025.06.22.660942

**Authors:** Rama D. Yammani, Xiaofei Chen, Nildris Cruz-Diaz, Xuewei Zhu, Swati Basu, Daniel B. Kim-Shapiro, David R. Soto-Pantoja, Leslie B. Poole

## Abstract

Anthracycline-induced cardiotoxicity remains a major limitation in cancer therapy, affecting long-term cardiovascular health in survivors. Dietary nitrate supplementation has shown cardioprotective effects in preclinical models of doxorubicin (Dox)-induced and ischemia-reperfusion injury. In this study, we evaluated the protective effects of nitrate and nitrite (NOx) supplementation in a syngeneic, tumor-bearing mouse model undergoing Dox chemotherapy (N=5 per group). NOx supplementation significantly reduced cardiac fibrosis and levels of 4-hydroxynonenal-modified proteins, a marker of oxidative stress. Tumor sizes varied, but most regressed by the final Dox dose. Importantly, NOx did not compromise the anti-tumor efficacy of Dox nor did it promote pulmonary metastasis. These findings support the potential clinical use of dietary nitrate and nitrite as adjuncts to Dox treatment to mitigate cardiotoxicity without impairing anti-cancer outcomes.

**Graphical Abstract:** 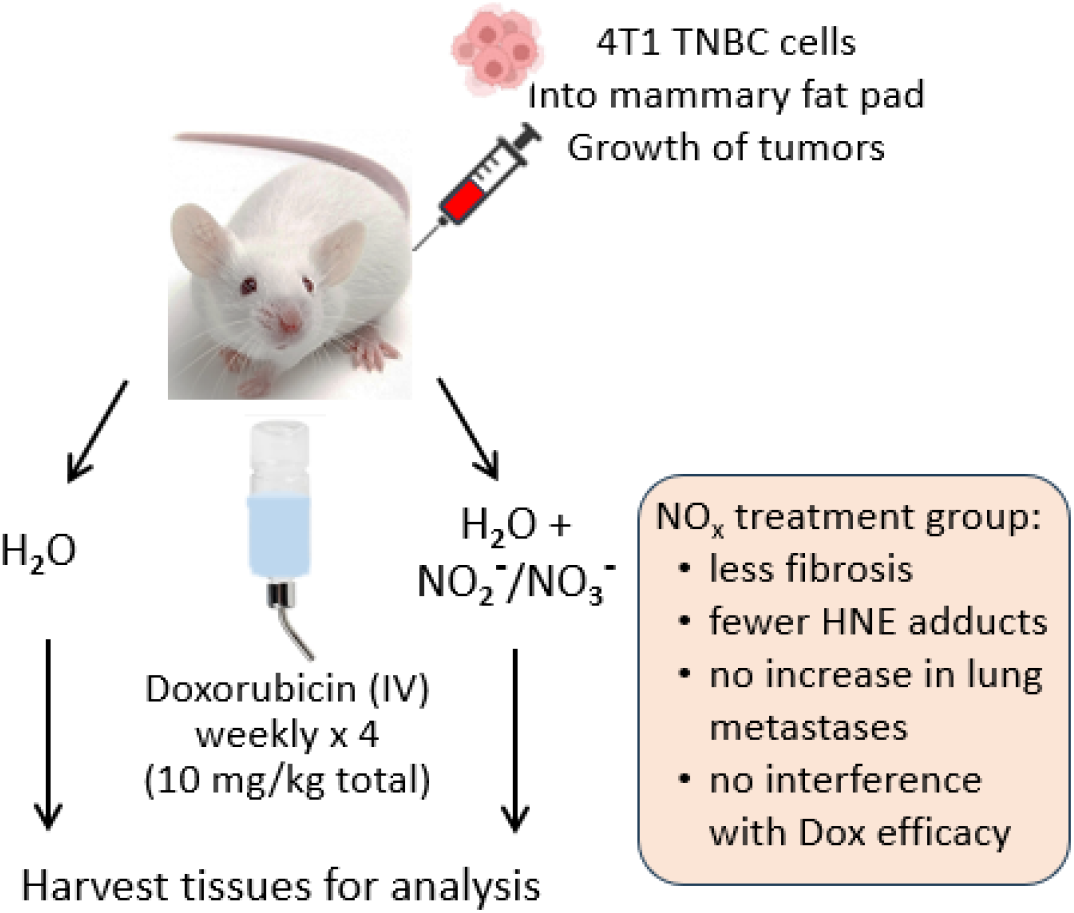

## 1. Introduction

Cardiotoxicity as a result of anthracycline chemotherapy is a significant health concern for large populations of survivors. Anthracycline administration is widespread as these are potent drugs employed, for example, against especially hard-to-treat, triple-negative breast cancer (TNBC) for which targeted therapies are ineffective. Unfortunately, their off-target toxic effects on heart function, both acute and chronic, are significant and dose-limiting. The urgency of this problem is driving extensive research in this area.

A number of cardioprotective co-therapies being tested focus on oxidative stress and redox-mediated protection, as anthracyclines like doxorubicin (Dox) generate reactive oxygen species (ROS), including peroxynitrite, particularly in mitochondria [1, 2]. To date, one cardioprotective therapy, an iron chelator (dexrazoxane), is approved for clinical use, but is rarely recommended due to problematic side effects [2]. Potential new therapies continue to be tested, including dietary nitrate and hydrogen sulfide which are cardioprotective in rodent models [3, 4], but much more research is needed to validate these new therapies and understand both the toxic and protective mechanisms involved.

One approach, as developed recently by members of our research group, is to generate reactive oxygen species (ROS)-triggered prodrugs that release both the anthracycline (e.g. doxorubicin, or Dox) and hydrogen sulfide, with the goal of maintaining or enhancing Dox efficacy against the tumor while protecting the heart and vasculature from drug-mediated damage [5], but this awaits further testing. Another approach focuses on dietary nitrate provided before and/or during Dox administration, as ingested nitrate is well established to be actively taken up by the salivary glands and provide positive effects (protection from ischemia-reperfusion injury to the heart, and enhancement of exercise efficacy) over sustained periods [3, 6-9]. Although molecular mechanisms of cardiac and vascular protection by dietary nitrate are not fully understood, multiple nitrogen oxide species (notably nitrite, NO _2_^**−**^; nitric oxide, ^•^NO and nitroxyl, HNO) are likely to play a role in these effects. Oral bacteria and other factors bioactivate dietary nitrate present in saliva to generate nitrite and ^•^ NO in blood and tissues, particularly under hypoxic conditions [6, 10]. Inhalation of sodium nitrite directly by patients has itself been shown to be of therapeutic value in combatting pulmonary arterial hypertension [11].

One study with mice challenged acutely with high-dose Dox demonstrated the efficacy of dietary nitrate in cardioprotection as shown by lowered cardiac fibrosis and lipid oxidation [3]. Still, to our knowledge, tumor-bearing animal models have not been tested. In the acute study, mice were sacrificed one week after challenge with a single high dose of Dox (15 mg/kg), testing whether sodium nitrate in the drinking water of adult male CF-1 mice was cardioprotective. Data obtained showed that the nitrate significantly decreased the Dox-induced impairment of ventricular function, decreased tissue lipid peroxidation, and protected against altered levels of specific antioxidant proteins in mitochondria (shown in a subsequent paper conducting proteomics analyses on these samples) [12]. Mechanisms through which the nitrate is cardioprotective are only poorly defined, but proteomics of cardiac tissue indicated significant increases, with Dox exposure, in the levels of the mitochondrial peroxiredoxin protein, Prx3, which was not increased in the nitrate-treated animals [12]; protection of cardiomyoblasts by hydrogen sulfide *in vitro* was also linked to similar changes in Prx3 levels [13], although it is not clear how keeping Prx3 levels low would offer a benefit against Dox toxicity.

Acute models of the cardiotoxic effects of chemotherapies have utilized large doses and tumor-free animals, neither of which adequately reflects human exposures. With the present study, we specifically sought evidence for or against the hypothesis that the nitrate-Dox combination would maintain anti-cancer efficacy and perhaps even sensitize TNBC tumors treated with Dox. While animal numbers were limited to 5 mice per group for the two groups, with and without NOx supplementation, this key question needed to be addressed before moving forward with larger preclinical studies. Analyses of the tumor sizes and tissue samples from the Dox-treated mice supplied (or not) with nitrate and nitrite in their drinking water demonstrated that the supplemented animals exhibited less fibrosis and fewer 4-hydroxynonenal (HNE) adducts. Our data also suggested that nitrate and nitrite did not impact the oncologic efficacy of Dox to cause TNBC tumors to regress, nor did they promote metastasis to the lung (instead showing a trend toward decreasing metastasis). In sum, these results suggest that there would be significant value in pursuing more in-depth studies to evaluate the therapeutic values of NOx supplementation using additional animal models and human subjects.

## 2. Materials and Methods

### 2.1 Experimental design and treatment protocol

Female 6-week-old BALB/c mice (n = 10) were purchased from Jackson Laboratory and put on a low nitrate diet (Harlan Teklad TD 99366) [14] one day post arrival. One week later mice were each injected with syngeneic 4T1 triple negative breast cancer (TNBC) cells (1 × 10^6^ cells) in the left fourth mammary gland fat pad to induce tumors. Mice were randomly divided into two groups: the first (control) group received doxorubicin therapy (Sigma, D1515), and the second (test) group was continuously provided with both sodium nitrate (1 g/L) and sodium nitrite (0.9 g/L) in their drinking water (replaced weekly) in addition to the doxorubicin therapy, according to the schedule shown in **Figure 1**. Nitrate levels used in the supplemented water were based on an earlier acute Dox toxicity study with mice [15], and nitrite levels, providing shorter-term effects, were based on Raat et al. [14]

Once the tumor volume reached 100 mm^3^, 2.5 mg/kg of IV-doxorubicin was injected into the tail vein once weekly for 4 weeks to reach an accumulated dose of 10 mg/kg. Tumor volumes were measured twice per week and echocardiography was performed before the first and after the last Dox treatment. At the study endpoint one week after the final dose of Dox, mice were humanely euthanized and plasma, tumors, lungs, and heart were harvested for analysis. All protocols were approved by the Animal Care and Use Committee of the Wake Forest University School of Medicine (protocol #A17-172, approved Feb 21, 2018), and all procedures were performed in accordance with relevant guidelines and regulations.

### 2.2 Echocardiography

Transthoracic echocardiography was performed using a Vevo 2100 LAZR ultrasound system (FUJIFILM/VisualSonics, Inc.; Toronto, Canada) equipped with a 30 MHz linear array transducer. M-mode short-axis images were obtained to assess ejection fraction and fractional shortening.

### 2.3 Nitrite and nitrate levels in plasma

Blood collected at the time of sacrifice was centrifuged (2,500 x g) for 3 min at 4 °C, then plasma was aliquoted into tubes, placed on dry ice, and stored at −80 °C until analysis by chemiluminescence. Plasma nitrate (NO_3_^**−**^) and nitrite (NO_2_^**−**^) levels were measured using an HPLC-based Eicom NOx Analyzer, model ENO-20, according to the instructions of the manufacturer. For all measurements, standard curves were obtained and used for quantitative measurements.

### 2.4 Tissue Preparation and Analysis

Tumors, lungs, and 2 mm cross-sectional slices taken approximately 5 mm from the bottom of the heart were fixed in 4% paraformaldehyde for 24 h before embedding in paraffin. Embedded lung and heart tissues were cut into 5 μm-thick sections and stained with hematoxylin and eosin (H&E). Lung metastatic lesions were quantified from the H&E images. Heart sections were also stained with PicroSirius Red for collagen, indicating areas of fibrosis; a total of 12 representative images from each group were quantified.

In addition, small pieces of cardiac tissue were rapidly frozen in liquid nitrogen at the time of sacrifice and used to conduct Western blotting for HNE adducts on proteins, and for levels of peroxiredoxin-3 (Prx3) and glyceraldehyde-3-phosphate dehydrogenase (GAPDH). Briefly, 11 mg of frozen heart tissue was weighed out and gently processed into a single cell suspension in 50 µl of sterile phosphate-buffered saline using the soft end of the 1 ml syringe plunger, with samples stored at −20 °C until analysis. Levels of Prx3 (antibody from Abcam, ab73349), 4-Hydroxynonenal (Abcam, ab46545), and GAPDH (EMD Millipore, CB1001) were evaluated with 3 technical replicates each by loading 2.5 µL per sample on a 10% polyacrylamide gel for SDS-PAGE, followed by Western blot analysis. Detection of antigens used horseradish peroxidase-linked secondary antibodies (Anti-rabbit IgG from Cell Signaling, catalogue # 7074 for HNE and Prx3 Western blots and Anti-mouse IgG from Cell Signaling, catalogue # 7076 for GAPDH) and SuperSignal West Dura Extended Duration Substrate kit from Thermo Scientific. Images were collected with a KODAK imager and analyzed using ImageJ software.

### 2.5 Statistical Analysis

Data are presented as the mean ± standard error of the mean (SEM). Statistical differences were evaluated using a non-paired, two-sided Student’s t-test (GraphPad Prism 10 Software, San Diego, CA, USA). The criterion for statistical significance was set at p<0.05.

Tumor volumes were obtained from measurements of the longest perpendicular axes ((long axes) × (short axes)^2^)/2).

## 3. Results

Based on the need to test nitrate/nitrite (NOx) effects on Dox cardiotoxicity and efficacy in tumor-bearing animal models, we designed a focused study with Dox-treated BALB/c mice bearing syngeneic breast cancer tumors where one of the two groups continuously received NOx in their drinking water; this mode of NOx administration is similar to the previous acute toxicity study with non-tumor bearing male CF-1 mice [3]. To maximize the ability to discern NOx effects, mice were maintained on a low nitrate diet [14]. As shown in **Fig.1A**, syngeneic 4T1 TNBC cells were injected into one mammary fat pad per mouse (N=10), then one week later, half of the mice (N=5) were switched to the NOx-containing water. Nitrate is the source of sustained NOx supplementation (through accumulation in the salivary glands and oral bioactivation); nitrite was also included as a more immediately bioavailable NOx source. Once tumors were palpable (about 2 weeks after injection of the 4T1 cells), echocardiography was performed, then all mice received four weekly intravenous doses of Dox (2.5 mg/kg each). One week after the final dose of Dox, mice were again evaluated by echocardiography, then sacrificed to collect various tissues (including plasma, lung and heart tissue). Analyses of the plasma verified that the supplementation of the drinking water was effective in elevating plasma levels of both nitrate and nitrite (**Fig. 1B** and **Fig.1C**, and Table S1).

**Figure 1.**
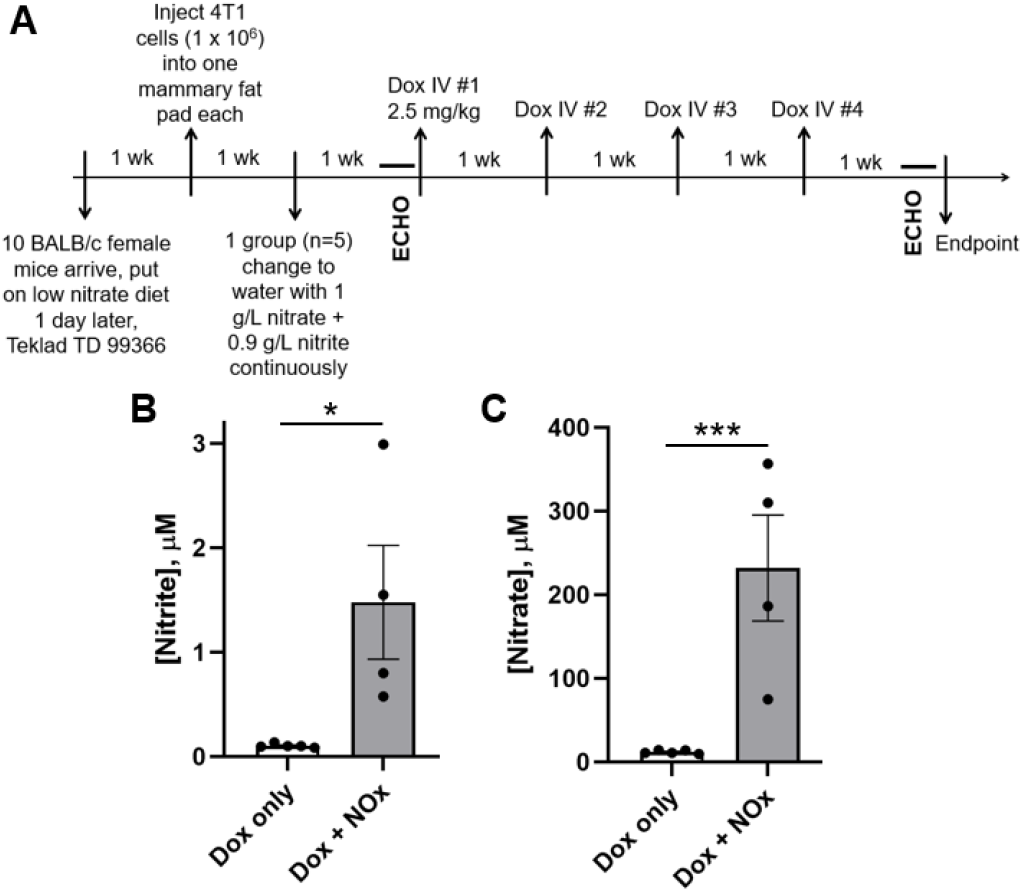
Experimental plan for assessing NOx supplementation effects on Dox-treated mice bearing triple negative breast cancer (TNBC) tumors, and associated nitrate and nitrite levels in plasma. **A**, Following injection of 4T1 cells and growth of tumors for 13 days, echocardiography (ECHO) was performed, then mice were treated with four weekly doses of Dox, again evaluated by ECHO, then sacrificed to harvest tissues for analysis. **B** and **C**, Plasma was evaluated for concentrations of nitrite (**B**) and nitrate (**C**). *, p<0.05, ***, p<0.001

As shown in **Fig. 2**, tumor volumes among the 10 mice varied, with one mouse never developing a tumor and another developing an unusually large tumor. Although the low animal numbers and variability limit the conclusions that can be made regarding any effect of NOx on Dox efficacy, our data suggest that dietary NOx supplementation does not impact Dox efficacy in this TNBC model.

**Figure 2.**
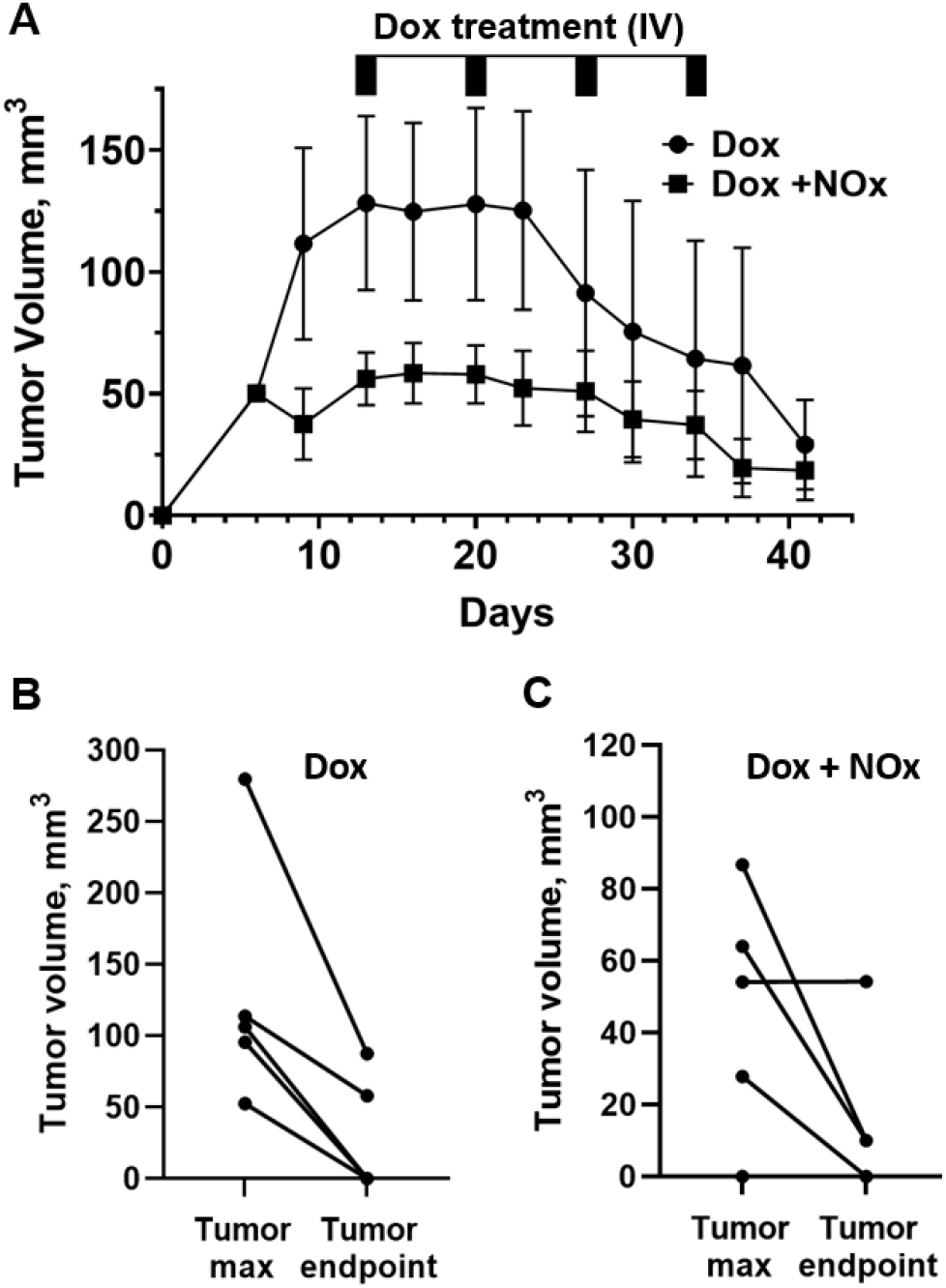
Tumor volumes of individual TNBC tumor-bearing BALB/c mice during Dox treatment in the presence or absence of NOx supplementation. **A**, Beginning 13 days after tumor cell injection, animals received four weekly intravenous (IV) doses of doxorubicin (2.5 mg/kg each), and tumor size was measured twice weekly using calipers. **B** and **C**, tumor volumes of mice treated with Dox alone (**B**), or Dox in combination with NOx-supplemented H_2_O (**C**), were compared at their maximal size and at the endpoint of 41 days.

While echocardiography data for both treatment groups did not reveal any Dox-mediated deficits (and therefore also no rescue by NOx treatments) as assessed by ejection fraction (EF) or fractional shortening (FS) (**Table S2**), a significant decrease in Dox-induced cardiac fibrosis assessed through PicroSirius Red staining of left ventricle (LV) slices was observed in the NOx-treated animals (**Fig. 3A** and **3B**). Moreover, oxidized lipid (4-hydroxynonenal, HNE) adducts on proteins detected by Western blots of LV tissue were suppressed by NOx treatment (**Fig. 3C**). In addition, the mitochondrial antioxidant protein Prx3, shown previously to be increased after acute Dox treatment [12], was lower in the mice treated with NOx (**Fig. 3D**).

**Figure 3.**
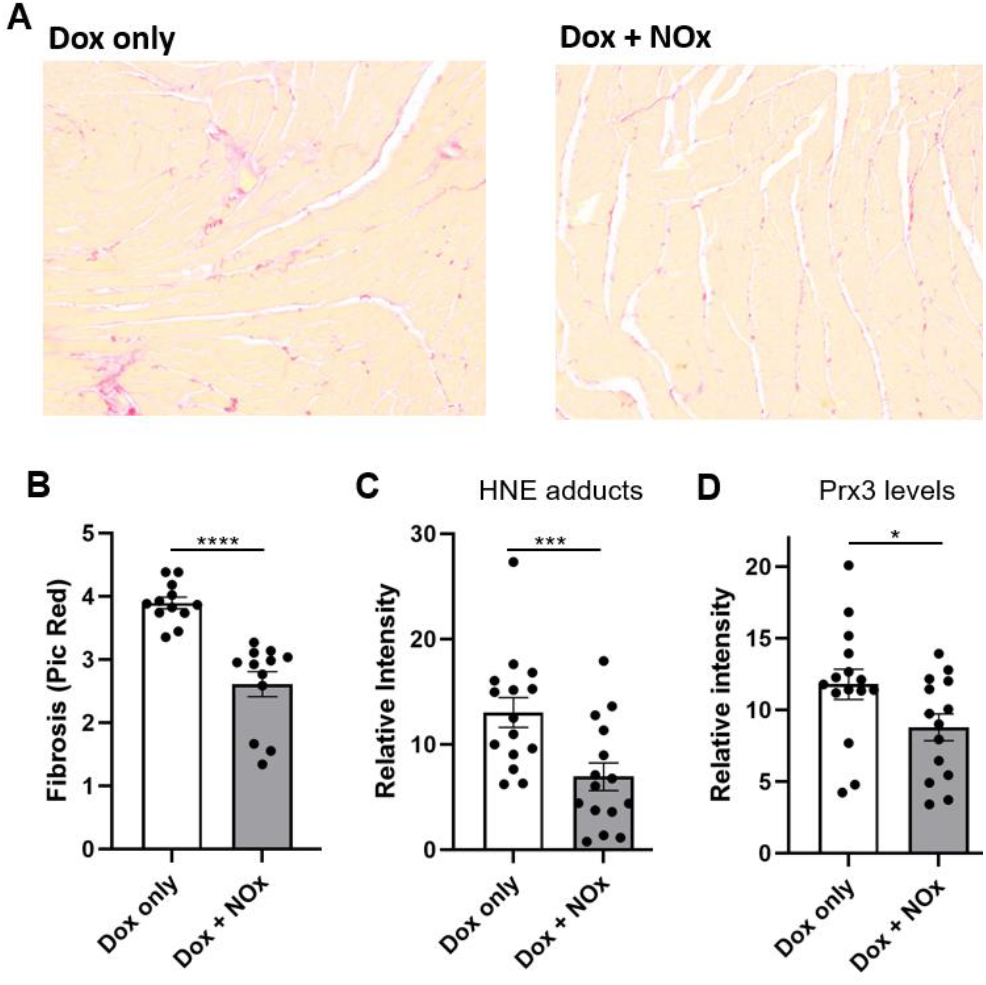
Protection of Dox-treated cardiac tissue from fibrosis and formation of protein adducts with 4-hydroxynonenal (HNE) by nitrate and nitrite supplementation. **A**, Fibrosis was evaluated using PicroSirius Red staining of left ventricle (LV) cross-sections. **B**, Fibrosis was significantly decreased with NOx treatment. **C** and **D**, HNE adducts **(C)**, and Prx levels **(D)**, were assessed by Western blot analysis with frozen LV tissue, indicating statistically-significant differences between groups. *, p<0.05, ***, p<0.001, ****, p<0.0001

Finally, metastatic lesions in lung tissue were quantified from slices of formalin-fixed lung tissue stained with Hematoxylin and Eosin. The lung slices were visually distinct between the NOx-treated and untreated animals, showing significant pathology in the absence of NOx (**Fig. 4A** and **4B**). Quantitation of metastatic lesions suggested fewer metastatic lesions in the NOx-treated mice (**Fig. 4C**). Still, this difference was not statistically significant (in part due to the overall low numbers of lesions as well as limited animal numbers).Importantly, there was no evidence that NOx supplementation caused an increase in lung metastases, which would be an outcome of concern.

**Figure 4.**
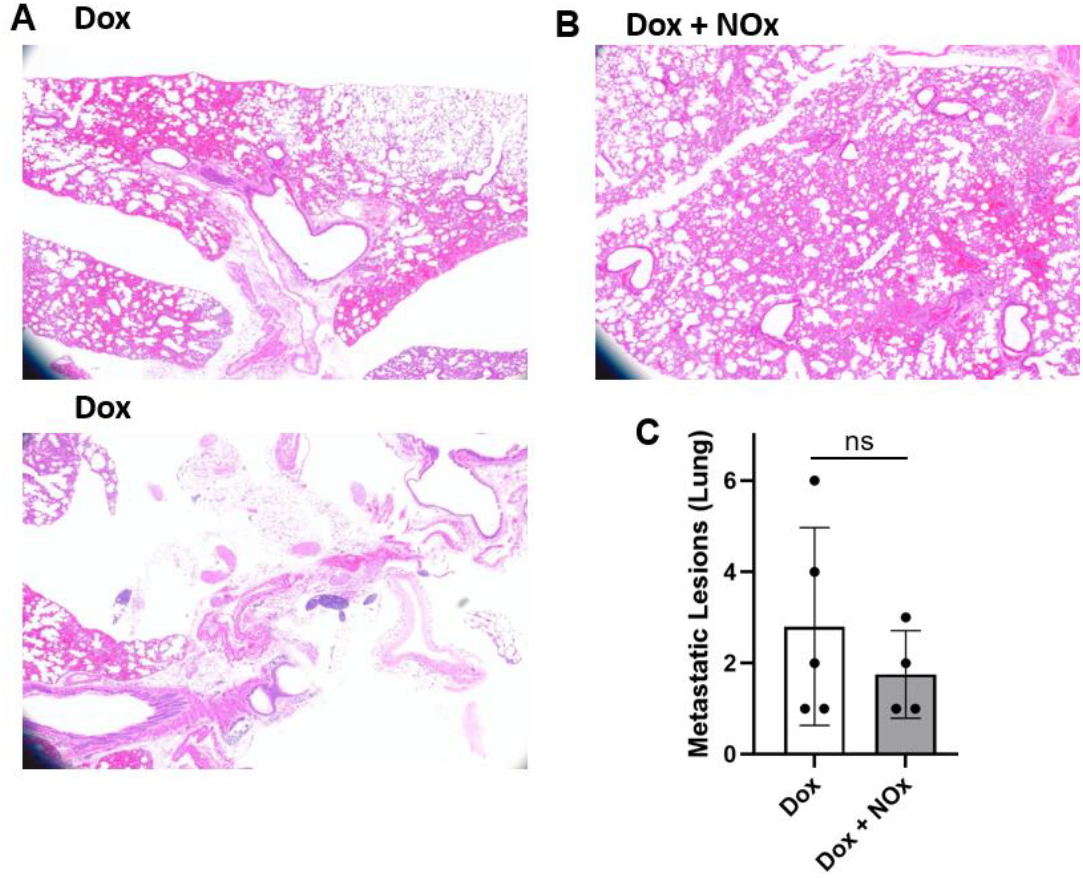
Effects of NOx supplementation on lung tissues and TNBC-associated metastatic lesions. **A** and **B**, Hematoxylin and Eosin (H&E) stained lung tissue from Dox-exposed mice showed significant evidence of tissue remodeling **(A)** which was less evident in the tissue from the NOx-supplemented mice **(B). C**, numbers of metastatic lesions in lung tissue associated with TNBC and Dox treatment were similar but trended lower in the NOx-supplemented animals (p=0.237). ns, not significant

## 4 Discussion

Multiple cardioprotective therapies are under investigation through preclinical or early clinical trials. Still, clinically-approved treatments to address cardiotoxicity of chemotherapeutics, other than dexrazoxane which is only approved for limited use, are not yet available. Inorganic nitrate (for example using beet root juice supplementation) holds promise as a cardioprotective agent given its demonstrated ability, in an acute Dox-exposure animal model, to preserve cardiac function and protect from oxidative damage [3]. In light of those findings in the absence of cancer, our team set out to determine whether the cardioprotective effect of nitrate would extend to TNBC tumor-bearing mice undergoing Dox-mediated therapy. We also sought preliminary indications of any adverse indications with respect to limitations on Dox efficacy or enhanced metastases to lung with nitrate supplementation, although a larger study will be needed to better address these issues. To ensure both short-term and long-term availability of NOx species, both NO _2_ ^**-**^and NO _3_^**-**^were included in the drinking water. While one animal in the NOx-treated group did not develop a tumor and tumor volumes were variable in the other mice, there was no indication that NOx supplementation impacted Dox anti-tumor activity (**Fig. 2A-C**). Analysis of the harvested tissues clearly indicated the protection, by NOx supplementation, of cardiac tissue from Dox-induced fibrosis (**Fig. 3A-B**) and oxidation-related protein damage as shown by HNE adducts detected by Western blot (**Fig. 3C**). Prx3 levels, which are known to rise with Dox treatment [12, 13], were also lower (**Fig. 3D**), perhaps due to the lower ROS levels in the NOx-protected animals. Functional studies using echocardiography (**Table S2**) were uninformative as there was no difference in ejection fraction (EF) or fractional shortening (FS) values with either Dox alone or in combination with NOx. It appears that protection of cardiac function by oral nitrate demonstrated previously in male CF-1 mice acutely exposed to Dox [3] was not detectable here in part because our chronic exposure model of breast cancer tumor-bearing female BALB/c mice exhibited lower EF and FS values at the outset that were not affected by Dox at the timepoints assessed (**Table S2**). Finally, and importantly, there was no evidence for an enhancement in metastatic TNBC lesions in the lungs of those animals supplemented with NOx; if anything, there was a trend toward fewer metastatic lesions in the NOx-treated animals. While preliminary, this study supports the need to study dietary nitrate and nitrite more deeply as cardioprotective agents compatible with anthracycline treatment. We postulate that nitrate, with or without added nitrite, may even lower the effective dose of Dox needed to treat the cancer; if true, this would further lower the risk of cardiovascular damage by anthracycline treatment. With its cardioprotective effects demonstrated here in a tumor-bearing mouse model, inorganic nitrate as an oral supplement could provide a cheap and convenient way to lower the risk of adverse cardiovascular effects of anthracycline treatment (as long as further confirmation is obtained that the anti-cancer effects of Dox are not impaired and the risk of metastatic spread of the cancer does not increase). Moreover, pre-existing hypertension in some patients exacerbates the cardiotoxic effects of chemotherapies, and as shown for pulmonary arterial hypertension noted above [11], NOx could also offer particular benefits for these patients.

It is well established that ROS generation plays a role in Dox cardiotoxicity, with involvement of (i) mitochondria-dependent production of ROS (in part due to accumulation of Dox and its binding to cardiolipin and redox cycling to produce superoxide in this organelle), (ii) ROS production from uncoupled (damaged) endothelial nitric oxide synthase (eNOS), and (iii) exacerbation of Dox cytotoxicity when cellular free iron levels are increased [2, 16, 17].

As yet, administration of simple antioxidants such as Vitamin E have not proven to be particularly effective, although other therapies continue to be explored for which lowering ROS is likely to contribute to the preliminary cardioprotective effects observed (e.g. with ACE inhibitors as well as NLRP3 inflammasome inhibitors like the flavonoid dihydromyricetin and resveratrol) [17, 18]. Dexrazoxane, initially utilized to counteract Dox toxicity based on its iron-chelating activity, was subsequently demonstrated to exert its cardioprotective effect primarily through its depletion of Topo2B, thereby limiting DNA damage and inflammatory responses [2]. It should be noted that ROS and reactive nitrogen species (RNS), and especially their interactions, are complex and highly dependent on context as their pleotropic effects, sometimes protective and sometimes damaging, are dependent on fluxes through multiple competing pathways (for example, accelerating superoxide removal by superoxide dismutase activity may suppress its reaction with nitric oxide to form peroxynitrite, but also enhances the production of hydrogen peroxide, another form of ROS). It is also important to recognize that simple chemical nitric oxide donors utilized in cell culture or animals do not exhibit the same properties as NOx *in vivo* [16], thus preclinical and clinical testing is imperative to test the effects of dietary nitrate administration properly.

There is an expanding appreciation for the nitrate – nitrite – ^•^ NO axis as an alternative way to generate protective ^•^NO, particularly as nitric oxide synthases cannot function without oxygen, which is a substrate. There is also a building recognition that the hypoxia-induced release of ^•^ NO from bioactivated nitrate in blood and other tissues is necessary for matching oxygen delivery to demand [10, 19]. Based on our work and that of others, it seems prudent to further pursue dietary inorganic nitrate as a protective agent against the insipid cardiotoxicity associated in many patients with chemotherapeutic treatments that effectively treat cancers but create a large survivor population with a potentially avoidable, long-persisting elevation in risk of cardiac failure.

## Acknowledgments

The authors thank Greg Hundley, Erik Barton, David Caudell, Nancy Kock, Paul Listrani, Nikolai Jenssen, Robert Wieland, Tom Forshaw, Adam Wilson and Jessica VonCannon for important contributions to study design, data collection and interpretation of results. Jasmina Varagic was also instrumental in this research which was conducted as part of her role as a faculty member in the Department of Surgery at Wake Forest University School of Medicine. This research was supported in part by pilot grants from the Center for Redox Biology and Medicine and the Comprehensive Cancer Center (NIH/NCI P30 CA012197) at Wake Forest University School of Medicine, as well as R01 GM119227 and R35 GM135179 to LBP, and American Cancer Society Research Scholar Grant RSG-19-150-01-LIB to DRS-P. Earlier studies that served as a pilot and informed the present experimental design were supported by a co-funded pilot grant from Wake Forest Innovations and the Clinical and Translational Science Institute at Wake Forest University Health Sciences (NIH/NCATS UL1 TR001420), and from the Center for Molecular Signaling of Wake Forest University.

## Author Contributions

Conceptualization, L.B.P.; experimental data collection, R.D.Y., X.C., N.C.-D., S.B.; data analysis, R.D.Y., D.B.K.-S., D.R.S.-P., L.B.P.; writing—original, review, and editing, R.D.Y., X.Z., D.B.K.-S., D.R.S.-P. and L.B.P. All authors have read and approved the final manuscript.

### Data Availability

Additional data not shown in the figures are reported in the Supplementary Material, including Table S1 showing plasma nitrate and nitrite values, and Table S2 showing echocardiography results.

## References

1. Mukhopadhyay, P., Rajesh, M., Batkai, S., Kashiwaya, Y., Hasko, G., Liaudet, L., Szabo, C. & Pacher, P. (2009) Role of superoxide, nitric oxide, and peroxynitrite in doxorubicin-induced cell death in vivo and in vitro, Am J Physiol Heart Circ Physiol. 296, H1466–83.

2. Hutchins, E., Yang, E. H. & Stein-Merlob, A. F. (2024) Inflammation in Chemotherapy-Induced Cardiotoxicity, Curr Cardiol Rep.

3. Zhu, S. G., Kukreja, R. C., Das, A., Chen, Q., Lesnefsky, E. J. & Xi, L. (2011) Dietary nitrate supplementation protects against Doxorubicin-induced cardiomyopathy by improving mitochondrial function, Journal of the American College of Cardiology. 57, 2181–9.

4. Zhang, H., Pan, J., Huang, S., Chen, X., Chang, A. C. Y., Wang, C., Zhang, J. & Zhang, H. (2024) Hydrogen sulfide protects cardiomyocytes from doxorubicin-induced ferroptosis through the SLC7A11/GSH/GPx4 pathway by Keap1 S-sulfhydration and Nrf2 activation, Redox biology. 70, 103066.

5. Hu, Q., Yammani, R. D., Brown-Harding, H., Soto-Pantoja, D. R., Poole, L. B. & Lukesh, J. C., 3rd (2022) Mitigation of doxorubicin-induced cardiotoxicity with an H2O2-Activated, H2S-Donating hybrid prodrug, Redox biology. 53, 102338.

6. Lundberg, J. O. & Weitzberg, E. (2022) Nitric oxide signaling in health and disease, Cell. 185, 2853–2878.

7. Salloum, F. N., Sturz, G. R., Yin, C., Rehman, S., Hoke, N. N., Kukreja, R. C. & Xi, L. (2015) Beetroot juice reduces infarct size and improves cardiac function following ischemia-reperfusion injury: Possible involvement of endogenous H2S, Experimental biology and medicine (Maywood, NJ. 240, 669–81.

8. Szponar, J., Nizinski, P., Dudka, J., Kasprzak-Drozd, K. & Oniszczuk, A. (2024) Natural Products for Preventing and Managing Anthracycline-Induced Cardiotoxicity: A Comprehensive Review, Cells. 13.

9. Kim, D. J., Gao, Z., Luck, J. C., Brandt, K., Miller, A. J., Kim-Shapiro, D., Basu, S., Leuenberger, U., Gardner, A. W., Muller, M. D. & Proctor, D. N. (2024) Effects of short-term dietary nitrate supplementation on exercise and coronary blood flow responses in patients with peripheral artery disease, Front Nutr. 11, 1398108.

10. Cortese-Krott, M. M. (2022) NO, nitrites and nitrates and their role in vascular control - New fundamental developments in basic and translational research in Hypertension News pp. 30–33

11. Zuckerbraun, B. S., George, P. & Gladwin, M. T. (2011) Nitrite in pulmonary arterial hypertension: therapeutic avenues in the setting of dysregulated arginine/nitric oxide synthase signalling, Cardiovascular research. 89, 542–52.

12. Xi, L., Zhu, S. G., Hobbs, D. C. & Kukreja, R. C. (2011) Identification of protein targets underlying dietary nitrate-induced protection against doxorubicin cardiotoxicity, J Cell Mol Med. 15, 2512–24.

13. Liu, M. H., Lin, X. L., Yuan, C., He, J., Tan, T. P., Wu, S. J., Yu, S., Chen, L., Liu, J., Tian, W., Chen, Y. D., Fu, H. Y., Li, J. & Zhang, Y. (2016) Hydrogen sulfide attenuates doxorubicin-induced cardiotoxicity by inhibiting the expression of peroxiredoxin III in H9c2 cells, Molecular medicine reports. 13, 367–72.

14. Raat, N. J., Noguchi, A. C., Liu, V. B., Raghavachari, N., Liu, D., Xu, X., Shiva, S., Munson, P. J. & Gladwin, M.T. (2009) Dietary nitrate and nitrite modulate blood and organ nitrite and the cellular ischemic stress response, Free Radic Biol Med. 47, 510–7.

15. Beyer, J. N., Hosseinzadeh, P., Gottfried-Lee, I., Van Fossen, E. M., Zhu, P., Bednar, R. M., Karplus, P. A., Mehl, R.A. & Cooley, R. B. (2020) Overcoming Near-Cognate Suppression in a Release Factor 1-Deficient Host with an Improved Nitro-Tyrosine tRNA Synthetase, J Mol Biol. 432, 4690–4704.

16. Daiber, A., Gori, T. & Munzel, T. (2011) Inorganic nitrate therapy improves Doxorubicin-induced cardiomyopathy a new window for an affordable cardiovascular therapy for everyone?, Journal of the American College of Cardiology. 57, 2190–3.

17. Octavia, Y., Tocchetti, C. G., Gabrielson, K. L., Janssens, S., Crijns, H. J. & Moens, A. L. (2012) Doxorubicin-induced cardiomyopathy: from molecular mechanisms to therapeutic strategies, Journal of molecular and cellular cardiology. 52, 1213–25.

18. Bjelogrlic, S. K., Radic, J., Jovic, V. & Radulovic, S. (2005) Activity of d,l-alpha-tocopherol (vitamin E) against cardiotoxicity induced by doxorubicin and doxorubicin with cyclophosphamide in mice, Basic Clin Pharmacol Toxicol. 97, 311–9.

19. Kuhn, V., Diederich, L., Keller, T. C. S. t., Kramer, C. M., Luckstadt, W., Panknin, C., Suvorava, T., Isakson, B. E., Kelm, M. & Cortese-Krott, M. M. (2017) Red Blood Cell Function and Dysfunction: Redox Regulation, Nitric Oxide Metabolism, Anemia, Antioxid Redox Signal. 26, 718–742.

