## Supplemental Tables 1 and 2 for "Dietary nitrate and nitrite protect against doxorubicin-induced cardiac fibrosis and oxidative protein damage in tumor-bearing mice"

**Supporting Information**

Table S1. Nitrate and nitrite levels in the plasma of NOx-treated and untreated mice.

| NOx species measured | Group 1 (Dox) | Group 2 (Dox with NOx) |
|---|---|---|
| Nitrite (μM) | 0.106 (± 0.008) | 1.48 (± 0.49) |
| Nitrate (μM) | 12.2 (± 0.9) | 239 (± 57) |

**Table S2.** Echocardiography results for ejection fraction and fractional shortening.

| Treatment, animal number | Ejection Fraction (%) | | Fractional shortening (%) | |
|---|---|---|---|---|
| | Start | End | Start | End |
| Dox only | | | | |
| **1751** | 64.3 | 61.0 | 34.6 | 32.0 |
| **1752** | 64.1 | 69.1 | 33.9 | 38.0 |
| **1753** | 61.0 | 61.0 | 31.7 | 31.8 |
| **1754** | 55.1 | 54.1 | 28.3 | 27.7 |
| **1755** | 66.5 | 65.1 | 36.0 | 35.0 |
| Dox + NOx | | | | |
| **1756** | 54.8 | 64.6 | 28.0 | 34.6 |
| **1757** | 50.7 | 57.7 | 25.5 | 29.9 |
| **1758** | 68.5 | 59.2 | 37.6 | 30.7 |
| **1759** | 69.6 | 57.5 | 38.4 | 29.7 |
| **1760** | 57.1 | 59.0 | 29.7 | 30.7 |
| Summary[1] | | | | |
| Dox only | 62.2 ± 2.0 | 62.1 ± 2.9 | 32.9 ± 1.3 | 32.9 ± 1.7 |
| Dox + NOx | 60.1 ± 3.8 | 59.6 ± 1.3 | 31.8 ± 2.6 | 31.1 ± 0.9 |

Mean ± standard error of the mean.
